# Protamine expression in somatic cells condenses chromatin and disrupts transcription without altering DNA methylation

**DOI:** 10.1101/2025.06.02.657337

**Authors:** Deepika Puri, Alexandra Bott, Monica Varona Baranda, Esra Dursun Torlak, Gina Esther Merges, Hubert Schorle, Wolfgang Wagner

## Abstract

Protamines play a crucial role in nuclear condensation during spermiogenesis, a process that involves significant chromatin remodeling and the replacement of histones. While much research has focused on the function of protamines in sperm development and fertility, their effects in non-sperm cells remain largely unexplored. In this study, we investigated the impact of overexpressing murine and human protamine 1 and 2 (PRM1 and PRM2) on nuclear architecture, histone eviction, DNA methylation, and transcription in HEK293T cells and mesenchymal stromal cells (MSCs). Overexpression of protamines resulted in nuclear condensation; particularly PRM1 showed notable enrichment in nucleoli, and cells exhibited cell cycle abnormalities. Immunofluorescence staining indicated a significant reduction in specific histone modifications (H3K9me3, H3K4me1, and H3K27Ac) in response to protamine expression, especially in MSCs. Interestingly, despite these changes in nuclear organization, the methylome remained largely stable. However, expression of protamines significantly diminished transcription, particularly of the ribosomal genes, upon PRM1 expression. Our studies indicate that PRM1 and PRM2 condense distinct genomic regions in somatic cells, resulting in widespread silencing of gene expression.

## Introduction

Protamines are nucleoproteins that facilitate the higher-order compaction of sperm chromatin to protect the genetic integrity of the paternal genome [1, 2]. Spermatids of all mammals express protamine 1 (PRM1), whereas rodents and primates additionally express protamine 2 (PRM2) [1], with their expression ratio being important for fertility [2]. Protamines participate in a cascade of events during spermiogenesis that result in histone replacement, a sharp decline in transcription, and nuclear condensation [2]. The chromatin structure is severely altered by the binding of protamines, predominantly to the major groove, neutralizing the phosphodiester backbone charge [1]. PRM2 is transcribed as a precursor; the N-terminal region of the translated PRM2 is cleaved during spermiogenesis, and the mature PRM2 associates with the condensing DNA [3]. During sperm development, protamines replace most histones, and this histone-to-protamine transition involves the replacement of somatic histones with testis-specific variants and the incorporation of transition proteins, followed by protamines [4]. Notably, a portion of histones and their modifications are retained and are thought to contribute to transmitting epigenetic memory from the sperm to the embryo [5].

The histone code is intertwined with the DNA methylation pattern [6, 7], and there is crosstalk between the two in development as well as diseases [8, 9]. Therefore, the histone-protamine transition may alter the methylome, but this has so far not been investigated. Analysis of global DNA methylation levels with either enzyme-linked immunosorbent assay or immunofluorescent analysis of 5-methylcytosine suggested that protamine deficiency in sperm is associated with increased global DNA methylation [10, 11]. Furthermore, sperm from patients with protamine deficiency showed higher DNA methylation at six of seven tested imprinted control genes [12], but it remains unclear if protamines contribute directly to this change in DNA methylation. Additionally, given that protamination leads to condensation of sperm DNA up to six times more than during a mitotic cycle, it is evident that access to the transcriptional machinery is severely impaired, resulting in near-complete silencing of transcription in sperm [13].

While the nuclear compaction during spermiogenesis has been well studied, there is little understanding of the effect of protamine expression in somatic cells. Previous studies have shown partial condensation of sheep fibroblast nuclei upon overexpression of PRM1; condensation of HEK293T nuclei upon overexpression of PRM2, and reduced proliferation of HeLa cells upon protamine overexpression [14-16]. To understand how PRM1 or PRM2 influences transcription, histone modifications, or DNA methylation in somatic cells, we have comprehensively analyzed the effect of overexpressing human or murine protamines in different cell types. Our results shed light on the effect of protamines in somatic cells and emphasize the distinct function of these proteins in chromatin condensation.

## Experimental Procedures

### Cell culture

The HEK293T cells were cultured in Dulbecco’s Modified Eagle Medium (DMEM), high glucose, supplemented with 10% fetal bovine serum (FBS) and 1% penicillin-streptomycin in an incubator at 37°C in a humidified atmosphere with 5% CO2. MSCs were isolated from human bone-marrow samples of patients undergoing orthopaedic surgery after written and informed consent (ethics approval EK300/13). The cells were cultured in DMEM low-glucose supplemented with pooled human platelet lysate (10%), L-glutamine (2 mM), penicillin– streptomycin (100 U ml−1) and heparin (5 IU ml−1) at 37 °C and 5% CO2. Mouse embryonic fibroblasts (MEFs) were prepared from WT 129s2/Sv mice as described before [1], and cultured in DMEM containing Glutamax, 4.5 g/l D-Glucose, and Pyruvate, 10% FBS, 1x essential amino acids, 1x non-essential amino acids, 1x L-Glutamine 50 µg/ml Penicillin/Streptomycin. The cell culture media were changed every 2-3 days, and the cells were passaged when the monolayer reached a confluency of 80-90%.

### Plasmids and transfection of cells

The plasmids pcDNA3.1-EGFP, pcDNA3.1-EGFP-PRM1 and pcDNA-3.1-EGFP-PRM2 were purchased from Genescript. The pEGFP-N3-Prm2 plasmid was generated as described before [2]. pEGFP-N3-Prm1 plasmid was constructed using the pEGFP_N3 (Clontech (#6080-1)) plasmid and *Prm1* amplified from C57Bl/6J mouse testis cDNA in the same manner and provided by the Schorle lab. The following overhang primers were used: Prm1_EcoRI_fwd (AAAAG AATTC ATGGC CAGAT ACCGAT G); Prm1_BamHI_rev (TTTTG GATCC GTATTT TTTAC ACCTT ATGGT G).

HEK293T cells were transfected with the plasmids, using the TransIT-LT1 transfection reagent (Mirus Bio). Three days post-transfection, HEK293T cells were sorted, and live, high EGFP-positive cells were used for further experiments. MSCs were transfected by electroporation using the NEON Transfection System using optimized protocols. 24 hours post-transfection, cells were selected with geneticin at 800µg/ml for 4 days. MEFs after passage 3 were transfected on 6-well plates at a confluency of approximately 50% in antibiotic-free MEF medium. The FUGENE HD Transfection reagent (Promega) was used according to the manufacturer’s instructions. Each well was transfected with 4 µg plasmid.

### Antibody staining and nuclear area measurement

Cells were stained with primary and secondary antibodies (Supplemental Table 2), followed by DAPI staining and imaged with a Zeiss Axio Observer fluorescence microscope, as described before [17]. FIJI [18] was used for the quantification of the nuclear area. Violin plots visualized the distribution of measured nuclear area values, and statistical significance was tested using the Mann-Whitney U test. The intensity of the histone modification staining was measured using FIJI using the mean integrated density measurements of the region of interest (ROI) identified based on DAPI staining. Statistical significance between conditions was tested using the Mann-Whitney U test.

### Quantitative reverse transcription Polymerase Chain Reaction (qRT-PCR)

Protamine transcription was analyzed by qRT-PCR in transfected HEK293T cells. Primer pairs were designed to target the human PRM1 and PRM2 genes and the mouse Prm1 and Prm2 genes (Supplemental Table 1). *GAPDH* was used as an internal control. The relative expression of the protamine samples was normalized to the relative expression of the empty vector samples to display the fold change.

### Apoptosis assay

Three days after transfection, HEK293T cells were collected and stained with Alexa Fluor 647 Annexin V solution and 5 µl of a 20 µg/ml DAPI solution following standard protocols and analyzed with the BD FACS Canto II. Stacked bar plots were generated to visualize the distribution across the four categories, and statistical significance between conditions was tested using Welch’s t-test.

### Cell cycle analysis

Three days after transfection, approximately 500,000 HEK293T cells per condition were stained with 1 µg/ml DAPI solution and transferred to FACS tubes. After 15 minutes of incubation, the cells were analyzed by flow cytometry with the BD FACS Canto II. Gates for the cell cycle phases were set on the DAPI histogram for each population. The cell cycle phases were analyzed in the high EGFP-positive population. Stacked bar plots were generated to visualize the distribution across the cell cycle phases, and statistical significance between conditions was tested using Welch’s t-test.

### DNA methylation analysis

DNA methylation profiles were analyzed in HEK293T cells transfected with empty vector, PRM1, or PRM2 using the Illumina Methylation EPIC v2.0 BeadChip. DNA from three replicate samples for each condition was isolated using the NucleoSpin Tissue kit (Macherey Nagel), and microarray analysis was performed at Life and Brain (Bonn, Germany). The R packages minfi (v.1.48.0) [19], SeSAMe (V 1.20.0) [20], limma [21], and missMethyl [22] were used for data import, preprocessing, quality control, and analysis, respectively. A multidimensional scaling (MDS) plot was created to visualize the overall similarity of methylation profiles across the different samples for the top 10,000 most variable CpG sites. Scatterplots of the mean beta values of the two conditions were created to visualize differentially methylated CpG sites (at least 20% difference of mean methylation).

### Transcriptomic analysis

Total RNA from HEK293T cells transfected with empty vector, PRM1 and PRM2 was isolated using the NucleoSpin RNA Plus kit. mRNA sequencing (mRNA-seq) with 3′-end enrichment was then performed using the QuantSeq 3′ mRNA Library Prep Kit (Lexogen) and sequencing on a NovaSeq 6000 platform. The mRNA-seq data was processed using the nf-core/rnaseq pipeline [23]. The pipeline involved quality control with FastQC, adapter trimming using Cutadapt, and alignment of reads to the genome using STAR [24]. Salmon was used for quantification of gene expression [25] and further normalization of the data was performed in R. After removal of mitochondrial genes, reads per kilo base of transcript per million mapped reads (RPKM) were calculated and further normalized by total number of reads per sample, as described in [26, 27]. Differentially expressed genes were computed using a t-test with p-values ≤ 0.05 adjusted by FDR. The feature Counts Biotypes tool was used to classify and quantify the reads based on the gene biotype annotations (e.g. protein-coding, pseudogene, lincRNA, etc.), by counting the number of reads that overlap with annotated gene features.

## Results

### Protamine expression leads to nuclear condensation

HEK293T cells were transfected with plasmids for expression of human and murine protamines as fusion proteins with enhanced green fluorescent protein (EGFP; Figure S1a, b), and EGFP high-expressing cells were sorted for further analysis. The expression of protamines was confirmed at the transcript level (Figure S1c). Whereas localization of EGFP in controls was rather cytoplasmic, the human PRM1/PRM2-EGFP fusion proteins and the murine Prm1/Prm2-EGFP fusion proteins exclusively localized to the nucleus (Figure 1a). Notably, the nuclear signal was not evenly distributed, but enriched in small nuclear areas that might correspond to nucleoli. Furthermore, protamine-expressing cells had significantly smaller nuclear areas compared to empty vector-transfected cells, suggesting nuclear condensation. Expression of PRM1, PRM2, Prm1, and Prm2 fusion proteins resulted in a reduction of mean nuclear area by 28.88%, 29.62%, 22.22%, and 31.11%, respectively (Figure 1b). When we transfected human mesenchymal stromal cells (MSCs) with PRM1 and PRM2, we observed the same significant reduction in mean nuclear size (22.72% and 34.71%, respectively; Figure 1c, d). The speckled nucleolar localization, especially for PRM1, was also observed in MSCs (Figure 1c). Furthermore, similar nuclear changes were observed in mouse embryonic fibroblasts (MEFS; Figure S1d), demonstrating that expression of protamines results in nuclear condensation in somatic cells.

**Figure 1.**
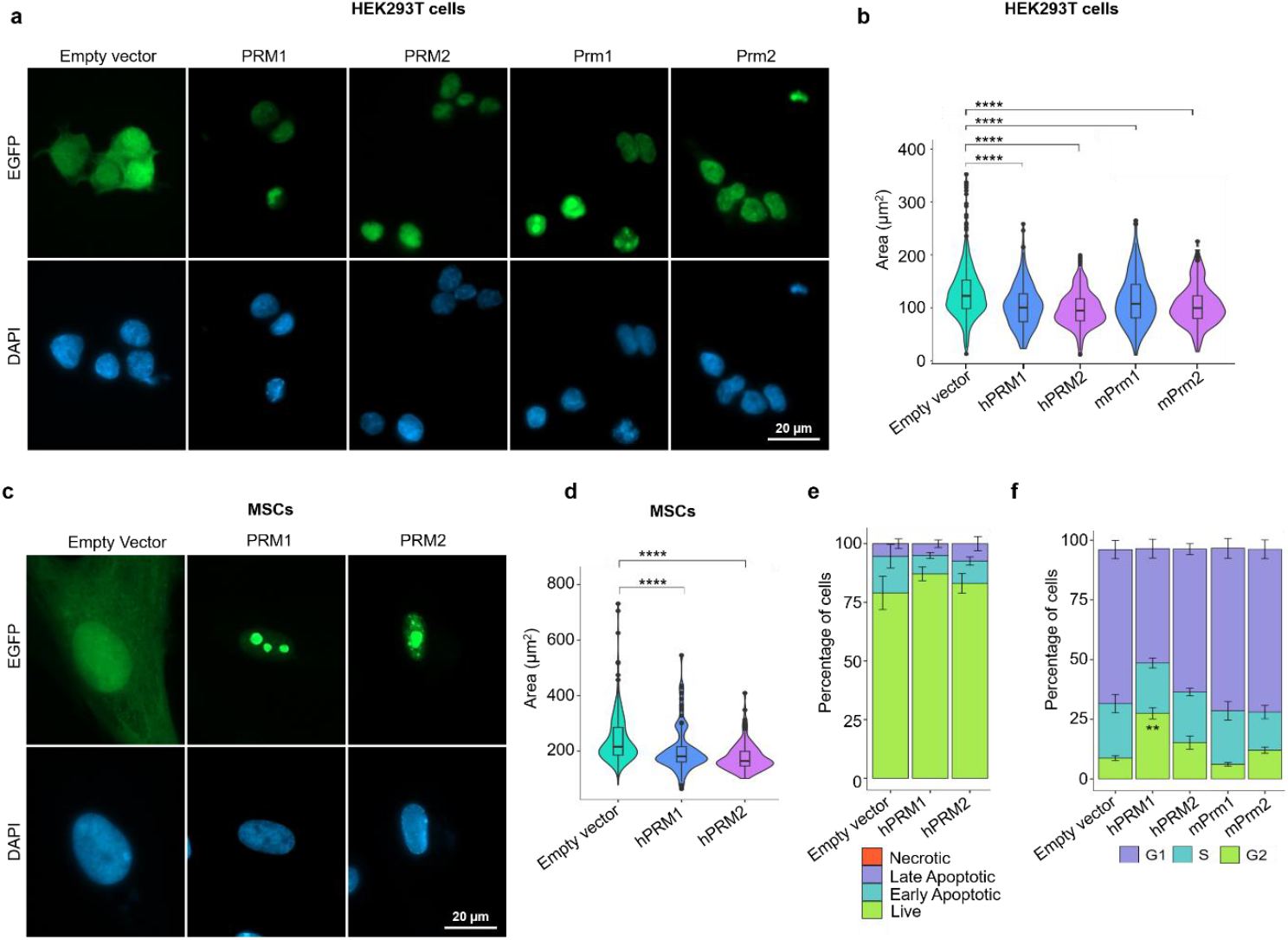
Protamine overexpression leads to nuclear condensation in somatic cells. **a)** Fluorescence microscopic analysis of HEK293T cells upon overexpression of protamine-EGFP fusion proteins or EGFP-control (nuclei are counterstained with DAPI). **b)** Nuclear area measurements in these cells. Violin plots indicate nuclear area (n = 3). P-values were calculated using the Welch’s t-test (****: p ≤ 0.00001). **c)** Fluorescence microscopic analysis of control and transfected MSCs stained with DAPI. **d)** Nuclear area measurements of MSCs transfected with empty vector or human protamine plasmids, in analogy to (a). **e)** Apoptosis analysis of control and transfected HEK293T cells (n = 3; error bars indicate the standard deviation). Welch’s t-tests did not reveal significant differences. **f)** Cell cycle analysis by flow cytometry of control HEK293T and cells transfected with human and murine protamines (n = 3; the error bars indicate the standard deviation). Significance was estimated with Welch’s t-test adjusted for multiple comparisons using the Benjamini-Hochberg method (**: p ≤ 0.001).

As nuclear condensation and nuclear membrane destabilization are characteristics of apoptosis [28], we performed Annexin PI staining. No significant differences were observed between the protamine and control conditions when comparing percentages of viable, early apoptotic, late apoptotic, and necrotic cells (Figure 1e), indicating that the nuclear condensation hardly resulted in apoptosis within 3 days of culture. The effects of protamine expression in HEK293T cells on cell cycle were further characterized by DAPI staining in flow cytometry. We observed a 2.6-fold higher percentage of cells in the G2/M phase and a 1.81-fold lower percentage of cells in the G1 phase in cells expressing PRM1 compared to empty vector cells. This effect was less in PRM2-expressing cells and not seen in murine Prm1 or Prm2-expressing cells (Figure 1f).

### Protamine expression diminishes histone modifications

Sperm protamination is associated with the near-complete eviction of histones during spermiogenesis [5]. To determine whether the nuclear condensation caused by protamine overexpression resulted in a displacement of histones, we quantified histone H3 along with histone modifications. Western blot analysis of global histone levels, H3K9me3, and H3K36me3 did not reveal obvious changes (Figure 2a). However, immunofluorescence staining showed a clear reduction of the H3K9me3 mark in HEK293T cells expressing PRM1 and PRM2 (Figure 2b, c). Overexpression of PRM1 in MSCs also showed a reduction in histone modification H3K9me3 (Figure 2d, e). Furthermore, in MSCs PRM1 and PRM2 expression resulted also in a marked reduction of H3K4me1 and H3K27Ac staining (Figure S2a, b). These results indicate that while the histones still persist, the nuclear localization of specific histone marks clearly diminishes upon protamine overexpression, indicating that there is partial histone displacement.

**Figure 2.**
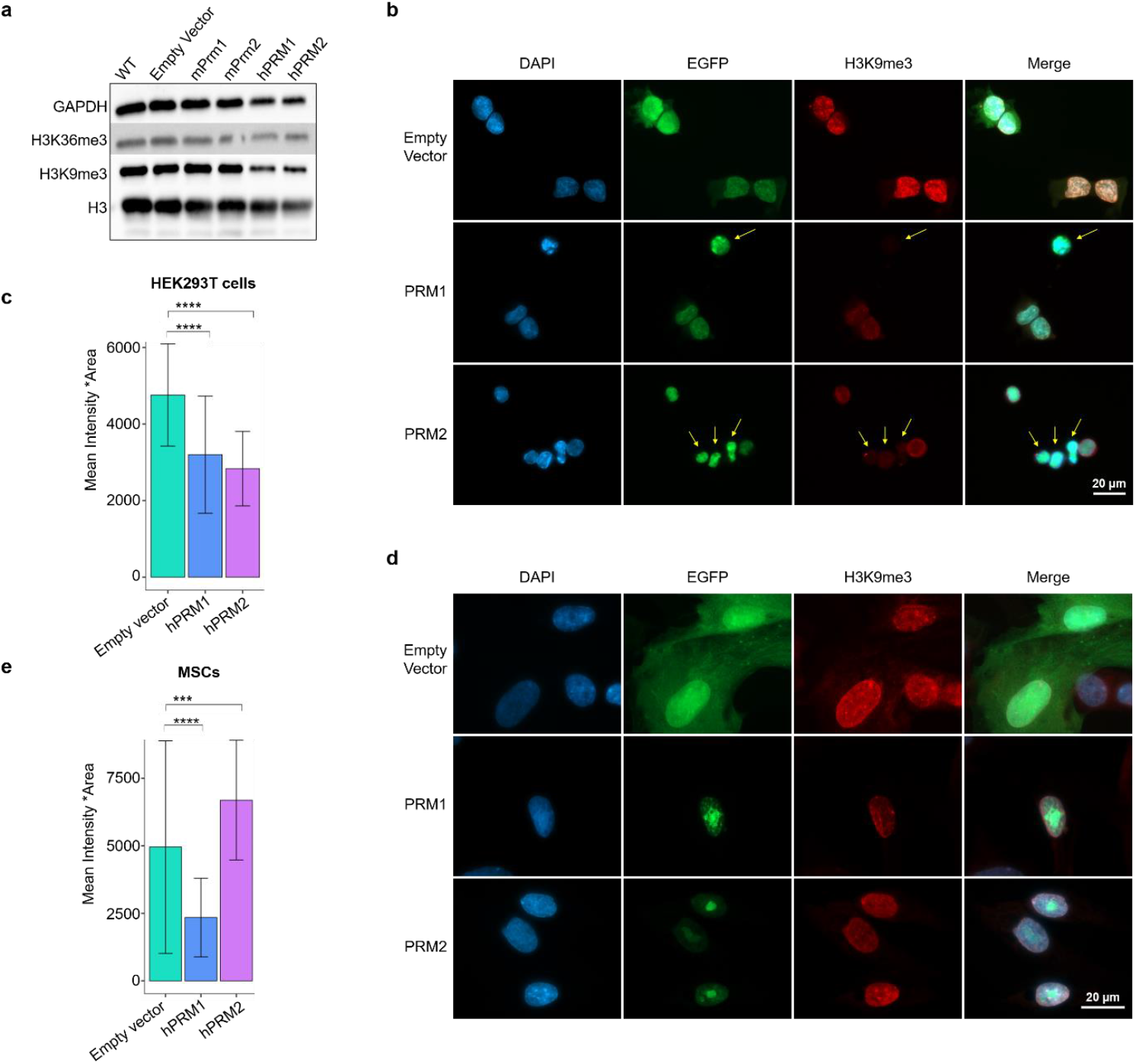
Protamine overexpression leads to displacement of histone modifications. **a)** Western blot analysis for histone H3, H3K36me3 and H3K9me3 in control and transfected HEK293T cells. GAPDH is used as a loading control. **b)** Fluorescence microscopy analysis of HEK293T cells transfected with control, PRM1, or PRM2 plasmids, and stained for H3K9me3 or DAPI. **c)** H3K9me3 intensity measurement of HEK293T cells (n = 3; mean intensity and standard deviation are indicated). P-values were calculated using the Welch’s t-test (**** : p ≤ 0.00001, ***: p ≤ 0.0001). **d**,**e)** Fluorescence microscopy analysis and H3K9me3 intensity measurement in MSCs, in analogy to (b and c).

### Protamine expression does not significantly affect DNA methylation

The nuclear condensation and partial histone displacement prompted us to investigate whether protamine expression also leads to changes in DNA methylation, as these epigenetic mechanisms are closely related [6, 8]. In fact, DNA methylation changes are very dynamic, occurring already within short timespans, as evidenced by the drastic DNA methylation changes in early development [29]. We therefore performed Illumina BeadChip analysis to determine DNA methylation profiles in controls and protamine-expressing HEK293T cells (n = 3). Principal component analysis (PCA) indicated that PRM1-expressing cells had heterogeneous DNA methylation patterns and clustered a bit apart from the controls and PRM2-expressing samples (Figure S3a). Notably, despite the clear nuclear condensation and histone displacement, none of the CG dinucleotides (CpGs) revealed significant differential methylation (limma adjusted P-value < .05). For orientation, we focused on the non-significant CpGs that showed at least 20% mean differential methylation. For PRM1, 617 CpGs were hypermethylated and 416 CpGs were hypomethylated; for PRM2, 38 CpGs were hypermethylated and 32 were hypomethylated (Figure 3a, b). There was a moderate overlap between differential DNA methylation of PRM1 and PRM2, which might also be due to the same empty vector controls utilized in both experiments (Figure 3c). Overall, nuclear condensation caused by protamine expression hardly resulted in reproducible changes in DNA methylation.

**Figure 3.**
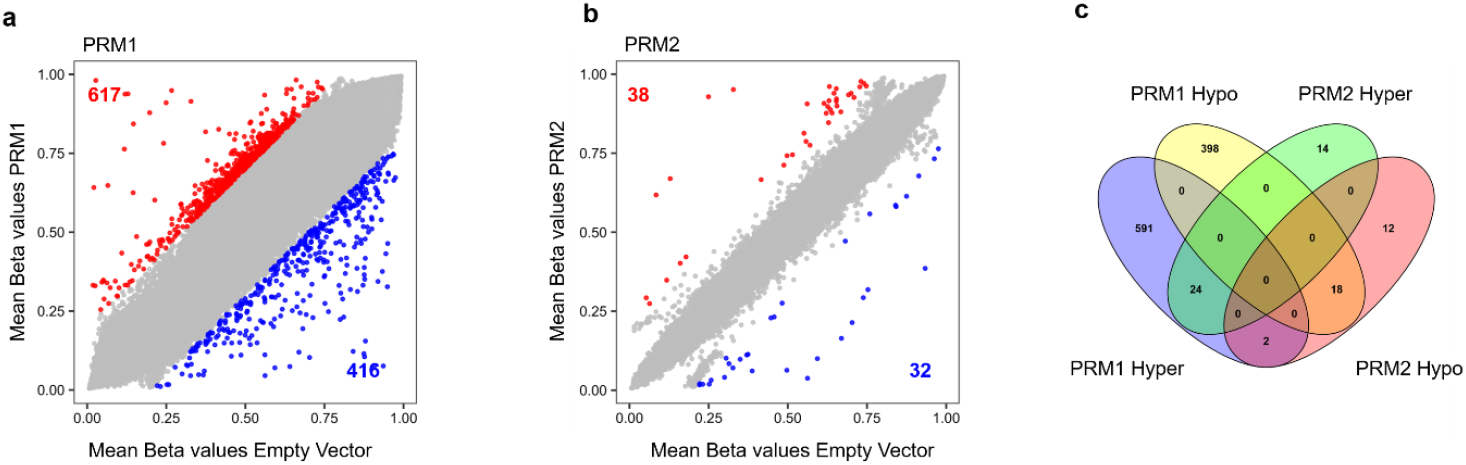
DNA methylome remains stable upon protamine overexpression. **a-b)** DNA methylation analysis of cells overexpressing PRM1 (a) or PRM2 (b) mean beta values, highlighting hypermethylated (red) and hypomethylated (blue) CpG sites (DiffMean cutoff = 0.2), but none of the CpGs reached statistical significance. **c)** Venn diagram depicting overlap of the in tendency differentially methylated CpGs upon overexpression of either PRM1 or PRM2.

### Protamine expression severely impairs transcription

During spermiogenesis, protamination is associated with an extensive reduction in transcription [30]; however, whether protamines directly cause transcriptional changes in somatic cells has not been explored. We performed RNAseq analysis of HEK293T cells expressing either PRM1 or PRM2, and found overall extensively reduced transcript levels in cells expressing protamines (Figure 4a). While an average of 23.4 million reads were obtained from control samples, only 10 million and 15.2 million reads were obtained from PRM1 and PRM2 expressing samples, respectively. Particularly, exonic reads decreased in PRM2-expressing cells, while PRM1-expressing cells exhibited similar percentages compared to empty vector transfection (Figure 4b). Furthermore, PRM2-expressing cells exhibited a higher representation of novel splice junctions and a reduction of known splice junctions and compared to empty vector and PRM1 (Figure 4c). This indicates that while overexpression of either PRM1 or PRM2 results in impaired transcription, there may be protamine-specific effects on the transcriptome.

**Figure 4.**
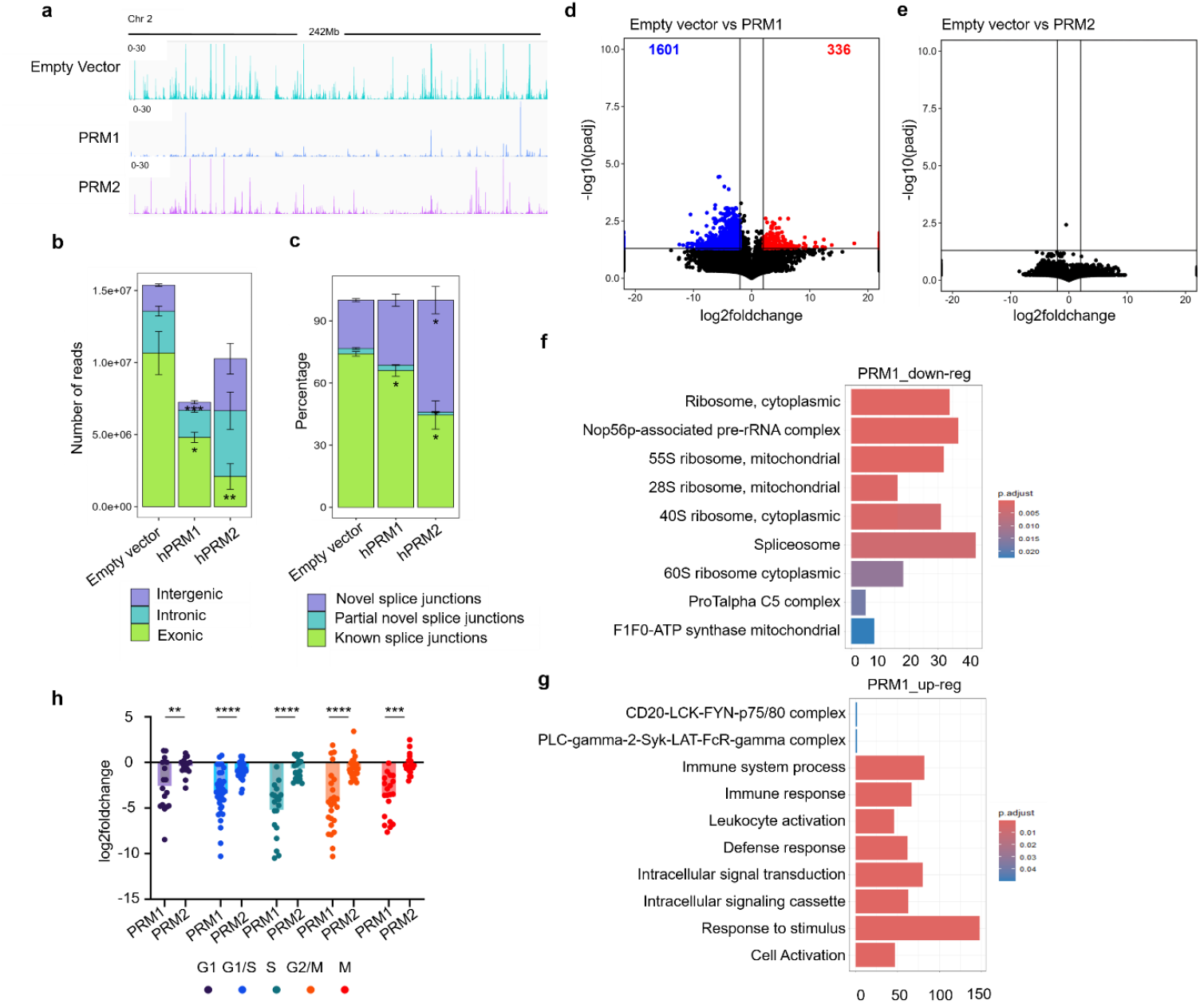
Protamine overexpression leads to severely impaired transcription. **a**) IGV tracks representing normalized transcription in control and transfected cells in a representative genomic region. **b)** Genomic origin of reads obtained by RNAseq in control and PRM1 and PRM2 transfected cells classified by Qualimap, and **c)** splice junctions classified by RSeQC (n = 3; mean and standard deviation are indicated). **d-e)** Volcano plots showing differential gene expression of upon overexpression of PRM1 (d) and PRM2 (e) compared to the empty vector controls. Significance thresholds (p≤ 0.05 and ≥ 2-fold change) are indicated by the dotted lines. **f-g)** Gene ontology analysis of upregulated (f) and downregulated (g) genes in PRM1-expressing HEK293T cells. **h)** Gene expression analysis of cell cycle genes in PRM1 and PRM2 expressing cells. The Y axis indicated log2fold change compared to the empty vector. Cell cycle genes are divided by the associated cell cycle phase. Significance was tested with the Welch’s t-test (**: p ≤ 0.001, ***: p ≤ 0.0001, ***: p ≤ 0.00001).

Due to the significantly lower overall level of transcription, the standard pipelines to determine differentially expressed genes (DEGs) may result in a normalization bias [26, 31]. We therefore performed DEG analysis after additional normalization by total number of reads per sample. Interestingly, we were only able to obtain DEGs for PRM1 overexpression, which resulted in 336 genes upregulated and 1601 genes downregulated, including DNA methyltransferase 3B (DNMT3B) and ten-eleven translocation methylcytosine dioxygenase 1 (TET1). In contrast, no significant differential expression was detected in PRM2-expressing cells. (Figure 4d, e), which may in part result from the high variability between the replicates of PRM2 expressing cells (Figure S3b), or a more random binding of PRM2 to accessible chromatin.

Gene ontology analysis of downregulated genes upon PRM1 overexpression showed significant enrichment of ribosomal, nucleolar, and splicing-related transcripts, which seem to preferentially include nucleolar transcripts (Figure 4f). This is in line with previous reports and our own data indicating nucleolar localization of PRM1 [14]. In contrast, upregulated genes were rather enriched in immune response and signalling pathway-related transcripts (Figure 4g).

Subsequently, we analyzed differential expression of cell cycle genes (KEGG pathway/ cell cycle/human) in PRM1 and PRM2 cells. While modest changes in cell cycle genes were seen in PRM2 cells, we observed a consistent reduction in cell cycle gene expression in PRM1 cells. This is particularly observed for S and G2 phase-associated genes (Figure 4h), which is consistent with our cell cycle analysis.

## Discussion

Our study demonstrates that overexpression of protamines in somatic cells results in a partial condensation of nuclei with disruption of transcription, while the methylome remains largely unaffected. Furthermore, PRM1 and PRM2 seem to have distinct effects on gene expression.

During spermiogenesis, protamines replace most histones while retaining histones and their modifications at developmentally important genomic locations [32, 33]. While we did not observe complete histone replacement upon expression of PRM1 or PRM2 in HEK293T or MSCs, it should be noted that the histone replacement during spermiogenesis is a part of a multistep cascade that begins with hyperacetylation of sperm-specific histones, incorporation of histone variants, nucleosome disassembly, DNA binding with transition proteins (TNPs) and finally replacement of the TNPs by protamines, each step sequentially facilitating nuclear compaction and transcription cessation [34]. Either way, we could observe a clear loss of H3K9me3 in HEK293T cells and MSCs expressing either PRM1 or PRM2. Previous studies demonstrated that transient expression of human or murine protamine 1 in sheep fibroblasts also resulted in loss of H3K9me3 [14, 35]. Furthermore, we saw either a reduction or altered localization of H3K4me1 and H3K27Ac marks upon PRM1 overexpression in MSCs, which might be attributed to the more open chromatin of MSCs as compared to HEK293T cells [36]. Previous studies have also demonstrated higher protamination and nuclear condensation in the presence of HDAC inhibitors such as trichostatin (TSA) [14, 35]. Hence, TSA might also enhance nuclear condensation upon protamine expression in somatic cells.

The DNA methylation pattern changes in a highly dynamic manner during cellular development and differentiation [29], and it has been shown that epigenetic crosstalk exists between different regulatory mechanisms [6, 7]. We therefore investigated whether nuclear reorganization with at least partial histone replacement could be reflected in the methylome. However, our results indicate that protamine expression does not evoke significant and directed changes in DNA methylation. In fact, previous studies compared DNA methylation levels in different stages of mouse spermatogenesis and saw a rather stable highly methylated profile throughout spermatogenesis [37]. Thus, the extensive nuclear condensation upon protamination might render the DNA inaccessible to DNA methylating or demethylating enzymes, resulting in a relatively stable methylome.

It is well known that cessation of transcription occurs during spermiogenesis [1, 13]. However, it is not understood whether and to what extent the protamines directly contribute to this transcriptional stop. Our results demonstrate that PRM1 and PRM2 expression severely impaired transcription, with a decline of 57.2% and 35% of the total transcripts, respectively. While the retained sequences showed similar distribution of exonic, intronic and intergenic regions for PRM1, PRM2 cells showed a significantly higher retention of intronic and intergenic sequences, which might point to different binding sites of these proteins. In fact, PRM2 is a zinc finger protein with a Cys2His2 motif [38], which may confer a DNA-binding and transcription regulation function distinct from PRM1 [39]. A report by Merges et al. revealed transcriptomic as well as proteomic differences of the testis of Prm1 knockout *versus* Prm2 knockout mice. Interestingly, GSEA showed enrichment of immune response genes in Prm1 - /-, but not Prm2 -/- mice, which is consistent with our results [40]. On the other hand, we observed clear down-regulation of ribosomal RNA genes upon PRM1 overexpression, and these genes are associated with nucleoli that also showed enriched EGFP signal in fluorescence microscopy. Thus, it appears likely that particularly PRM1 condenses the nucleolar genomic regions and thereby interferes more with ribosomal genes than the average silencing effect.

Taken together, protamines can alter chromatin structure and nuclear architecture even in somatic cells. Their property of nucleotide binding and chromatin condensation lends them to potential applications in nanopharmaceuticals and drug targeting [41, 42]. Protamines are considered to bind to DNA in a sequence-independent manner using electrostatic interactions between the DNA and the arginine-rich protamines [1]. This may preclude genomic regions associated with closed chromatin from being accessible to protamine binding. Our results indicate that PRM1 and PRM2 have different binding preferences. While both proteins lead to nuclear condensation, partial histone eviction and severely impaired transcription independent of DNA methylation, particularly the overexpression of PRM2 resulted in additional significant gene expression changes. It should be noted that the histone to protamine transition in human and mouse relies on a temporally regulated expression of both protamines, with protamine 2 expressed as a precursor, in a specific ratio [34]. While we haven’t explored this aspect of protamine incorporation in our study, further investigating these differences would not only elucidate better the process of sperm protamination but also assist in the development of potential protamine-mediated applications.

## Author contributions

D.P and W.W conceived the study. A.B performed the experiments involving transient transcription in HEK293T cells and MSCs under the supervision of D.P. G.E.M performed transient transfection in MEFs. M.V.B performed RNA seq analysis. E.D.T performed DNA methylation analysis. H.S provided the mouse Prm1 and 2 plasmids. D.P and W.W. wrote the manuscript. All authors read and approved the manuscript.

## References

1. Balhorn, R., The protamine family of sperm nuclear proteins. Genome Biol, 2007. 8(9): p. 227.

2. Arevalo, L., et al., Protamines: lessons learned from mouse models. Reproduction, 2022. 164(3): p. R57–R74.

3. Yelick, P.C., et al., Mouse protamine 2 is synthesized as a precursor whereas mouse protamine 1 is not. Mol Cell Biol, 1987. 7(6): p. 2173–9.

4. Wang, T., et al., Essential Role of Histone Replacement and Modifications in Male Fertility. Frontiers in Genetics, 2019. 10.

5. Torres-Flores, U. and A. Hernández-Hernández, The Interplay Between Replacement and Retention of Histones in the Sperm Genome. Frontiers in Genetics, 2020. 11.

6. Cedar, H. and Y. Bergman, Linking DNA methylation and histone modification: patterns and paradigms. Nat Rev Genet, 2009. 10(5): p. 295–304.

7. Rose, N.R. and R.J. Klose, Understanding the relationship between DNA methylation and histone lysine methylation. Biochimica Et Biophysica Acta-Gene Regulatory Mechanisms, 2014. 1839(12): p. 1362–1372.

8. Kondo, Y., Epigenetic cross-talk between DNA methylation and histone modifications in human cancers. Yonsei Med J, 2009. 50(4): p. 455–63.

9. de la Calle Mustienes, E., J.L. Gomez-Skarmeta, and O. Bogdanovic, Genome-wide epigenetic cross-talk between DNA methylation and H3K27me3 in zebrafish embryos. Genom Data, 2015. 6: p. 7–9.

10. Rajabi, H., et al., Pronuclear epigenetic modification of protamine deficient human sperm following injection into mouse oocytes. Systems Biology in Reproductive Medicine, 2016. 62(2): p. 125–132.

11. Rahiminia, T., et al., Relation between sperm protamine transcripts with global sperm DNA methylation and sperm DNA methyltransferases mRNA in men with severe sperm abnormalities. Human Fertility, 2021. 24(2): p. 105–111.

12. Hammoud, S.S., et al., Alterations in sperm DNA methylation patterns at imprinted loci in two classes of infertility. Fertility and Sterility, 2010. 94(5): p. 1728–1733.

13. Steger, K., Transcriptional and translational regulation of gene expression in haploid spermatids. Anat Embryol (Berl), 1999. 199(6): p. 471–87.

14. Iuso, D., et al., Exogenous Expression of Human Protamine 1 (hPrm1) Remodels Fibroblast Nuclei into Spermatid-like Structures. Cell Rep, 2015. 13(9): p. 1765–71.

15. Arevalo, L., et al., Loss of the cleaved-protamine 2 domain leads to incomplete histone-to-protamine exchange and infertility in mice. PLoS Genet, 2022. 18(6): p. e1010272.

16. Gunther, K., et al., Expression of sperm-specific protamines impairs bacterial and eukaryotic cell proliferation. Histochem Cell Biol, 2015. 143(6): p. 599–609.

17. Zeevaert, K., et al., YAP1 is essential for self-organized differentiation of pluripotent stem cells. Biomaterials Advances, 2023. 146.

18. Schindelin, J., et al., Fiji: an open-source platform for biological-image analysis. Nature Methods, 2012. 9(7): p. 676–682.

19. Aryee, M.J., et al., Minfi: a flexible and comprehensive Bioconductor package for the analysis of Infinium DNA methylation microarrays. Bioinformatics, 2014. 30(10): p. 1363–1369.

20. Zhou, W.D., et al., SeSAMe: reducing artifactual detection of DNA methylation by Infinium BeadChips in genomic deletions. Nucleic Acids Research, 2018. 46(20).

21. Ritchie, M.E., et al., limma powers differential expression analyses for RNA-sequencing and microarray studies. Nucleic Acids Res, 2015. 43(7): p. e47.

22. Phipson, B., J. Maksimovic, and A. Oshlack, missMethyl: an R package for analyzing data from Illumina’s HumanMethylation450 platform. Bioinformatics, 2016. 32(2): p. 286–288.

23. Ewels, P.A., et al., The nf-core framework for community-curated bioinformatics pipelines. Nat Biotechnol, 2020. 38(3): p. 276–278.

24. Dobin, A., et al., STAR: ultrafast universal RNA-seq aligner. Bioinformatics, 2013. 29(1): p. 15–21.

25. Patro, R., et al., Salmon provides fast and bias-aware quantification of transcript expression. Nature Methods, 2017. 14(4): p. 417-+.

26. Evans, C., J. Hardin, and D.M. Stoebel, Selecting between-sample RNA-Seq normalization methods from the perspective of their assumptions. Brief Bioinform, 2018. 19(5): p. 776–792.

27. Ignatov, D.V., et al., Dormant non-culturable Mycobacterium tuberculosis retains stable low-abundant mRNA. BMC Genomics, 2015. 16: p. 954.

28. Wyllie, A.H., et al., Chromatin Cleavage in Apoptosis -Association with Condensed Chromatin Morphology and Dependence on Macromolecular-Synthesis. Journal of Pathology, 1984. 142(1): p. 67–77.

29. Wilkinson, A.L., I. Zorzan, and P.J. Rugg-Gunn, Epigenetic regulation of early human embryo development. Cell Stem Cell, 2023. 30(12): p. 1569–1584.

30. Rathke, C., et al., Chromatin dynamics during spermiogenesis. Biochim Biophys Acta, 2014. 1839(3): p. 155–68.

31. Loven, J., et al., Revisiting global gene expression analysis. Cell, 2012. 151(3): p. 476–82.

32. Hammoud, S.S., et al., Distinctive chromatin in human sperm packages genes for embryo development. Nature, 2009. 460(7254): p. 473–8.

33. Puri, D., J. Dhawan, and R.K. Mishra, The paternal hidden agenda: Epigenetic inheritance through sperm chromatin. Epigenetics, 2010. 5(5): p. 386–91.

34. Moritz, L. and S.S. Hammoud, The Art of Packaging the Sperm Genome: Molecular and Structural Basis of the Histone-To-Protamine Exchange. Front Endocrinol (Lausanne), 2022. 13: p. 895502.

35. Palazzese, L., et al., Nuclear quiescence and histone hyper-acetylation jointly improve protamine-mediated nuclear remodeling in sheep fibroblasts. PLoS One, 2018. 13(3): p. e0193954.

36. Niwa, H., Open conformation chromatin and pluripotency. Genes Dev, 2007. 21(21): p. 2671–6.

37. Galan, C., et al., Stability of the cytosine methylome during post-testicular sperm maturation in mouse. PLoS Genet, 2021. 17(3): p. e1009416.

38. Bianchi, F., et al., P2 protamines from human sperm are zinc -finger proteins with one CYS2/HIS2 motif. Biochem Biophys Res Commun, 1992. 182(2): p. 540–7.

39. Razin, S.V., et al., Cys2His2 zinc finger protein family: classification, functions, and major members. Biochemistry (Mosc), 2012. 77(3): p. 217–26.

40. Merges, G.E., et al., Loss of Prm1 leads to defective chromatin protamination, impaired PRM2 processing, reduced sperm motility and subfertility in male mice. Development, 2022. 149(12).

41. Scheicher, B., A.L. Schachner-Nedherer, and A. Zimmer, Protamine-oligonucleotide-nanoparticles: Recent advances in drug delivery and drug targeting. Eur J Pharm Sci, 2015. 75: p. 54–9.

42. Ruseska, I., et al., Use of Protamine in Nanopharmaceuticals-A Review. Nanomaterials (Basel), 2021. 11(6).

