## Supplemental material for "Protamine expression in somatic cells condenses chromatin and disrupts transcription without altering DNA methylation"

### Protamines condense chromatin and disrupt transcription in somatic cells

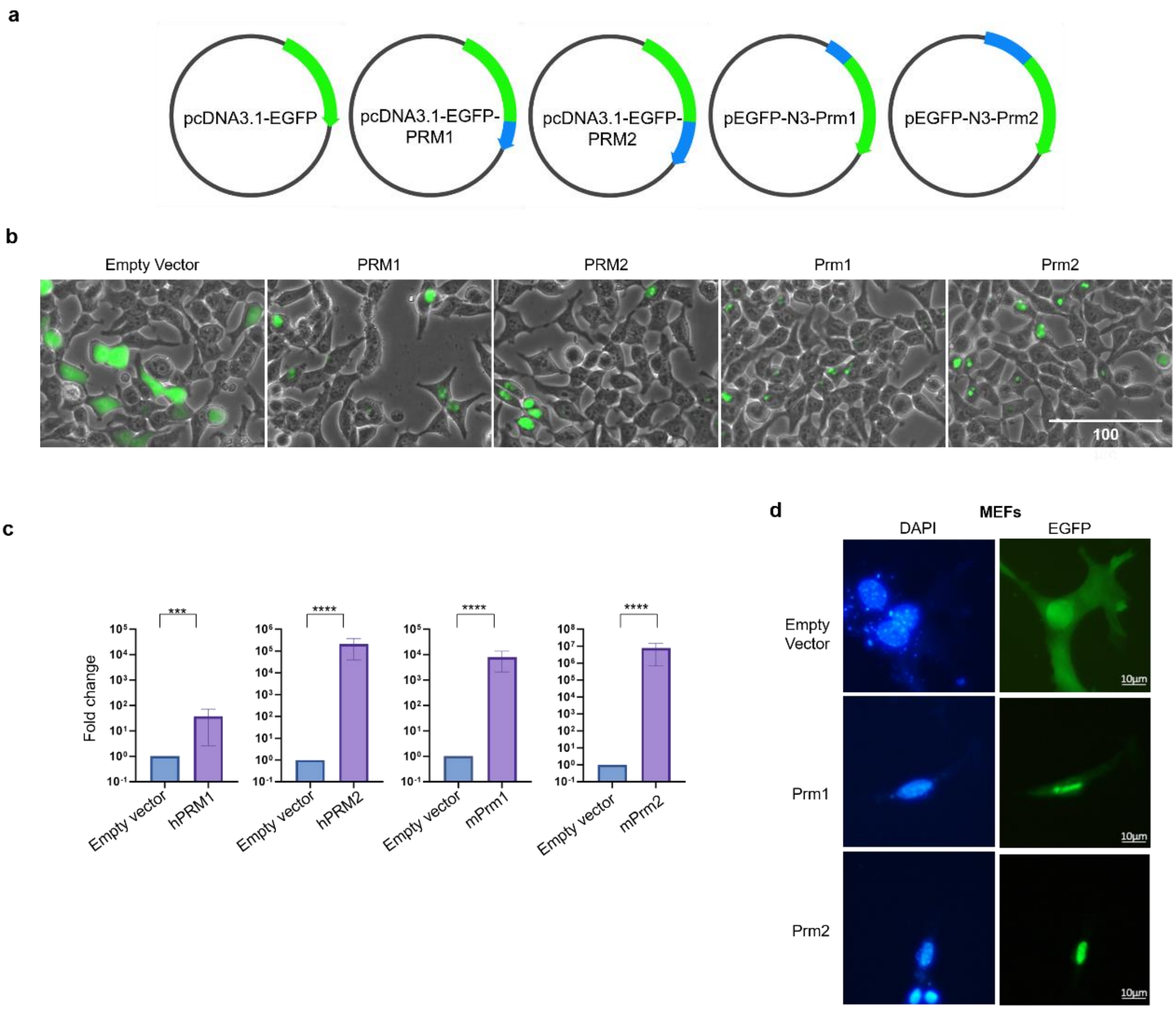


**Figure S1: Protamine overexpression and effects on somatic cells**
a) Schemes of plasmids. b) Brightfield images of HEK293T cells transfected with empty vector and human and murine protamines. c) qRT-PCR in HEK293T cells transfected with the plasmids pEGFP-empty, pEGFP-PRM1, pEGFP-PRM2, pEGFP-Prm1, and pEGFP-Prm2 (n = 3; fold change normalized to empty vector samples; error bars indicate standard deviation). P-values were calculated using the Welch’s t-test and adjusted for multiple comparisons using the Benjamini-Hochberg method (***: $p \leq 0.0001$, ****: $p \leq 0.00001$). d) Immunostaining analysis of control and transfected MEF cells stained with DAPI.

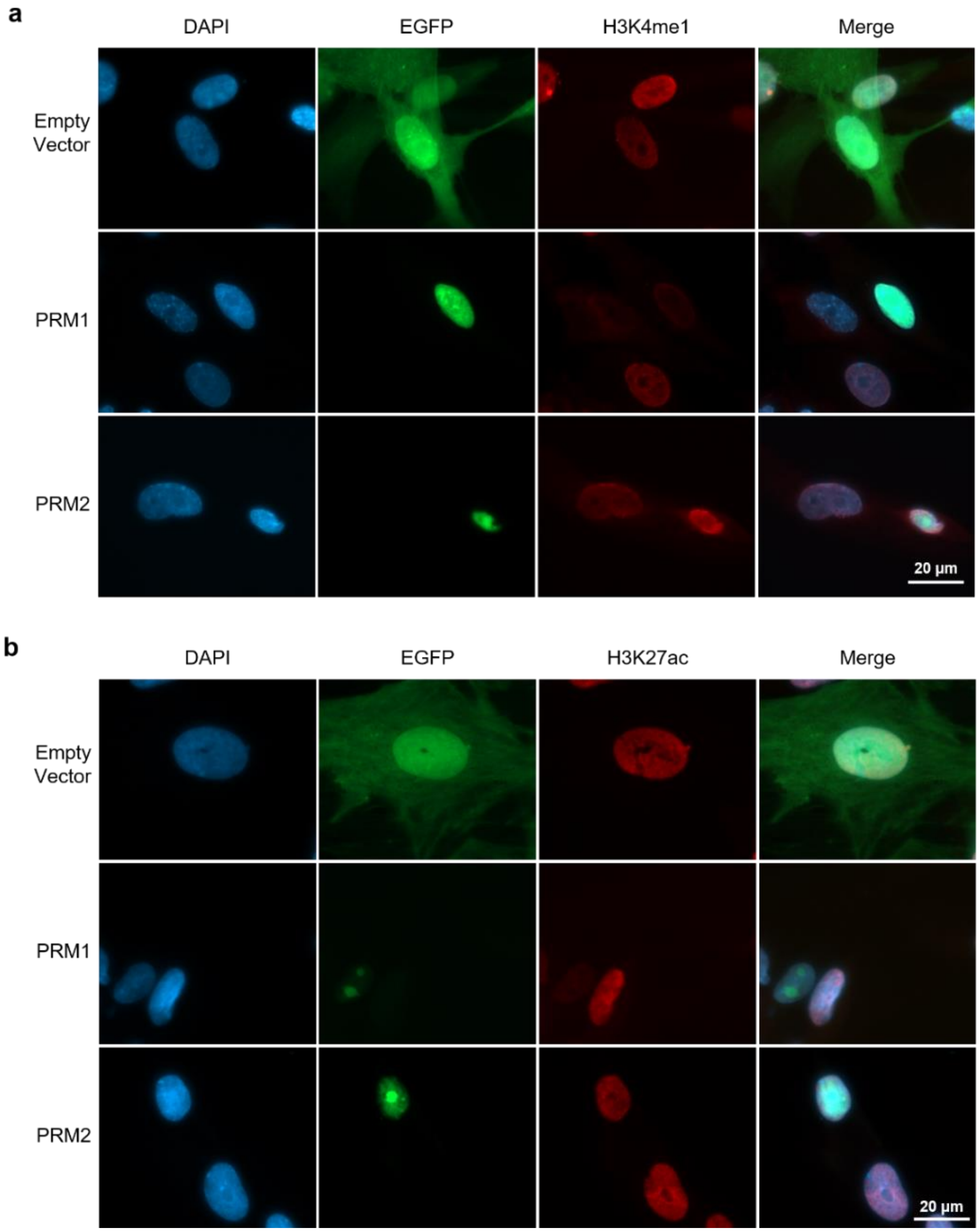


**Figure S2: Protamine overexpression displaces H3K4me1 and H3K27ac**
a-b) Fluorescence microscopy analysis of MSCs transfected with control, PRM1, or PRM2 plasmids, stained with for H3K4me1(a) and H3K27Ac (b) antibody and counterstained with DAPI.

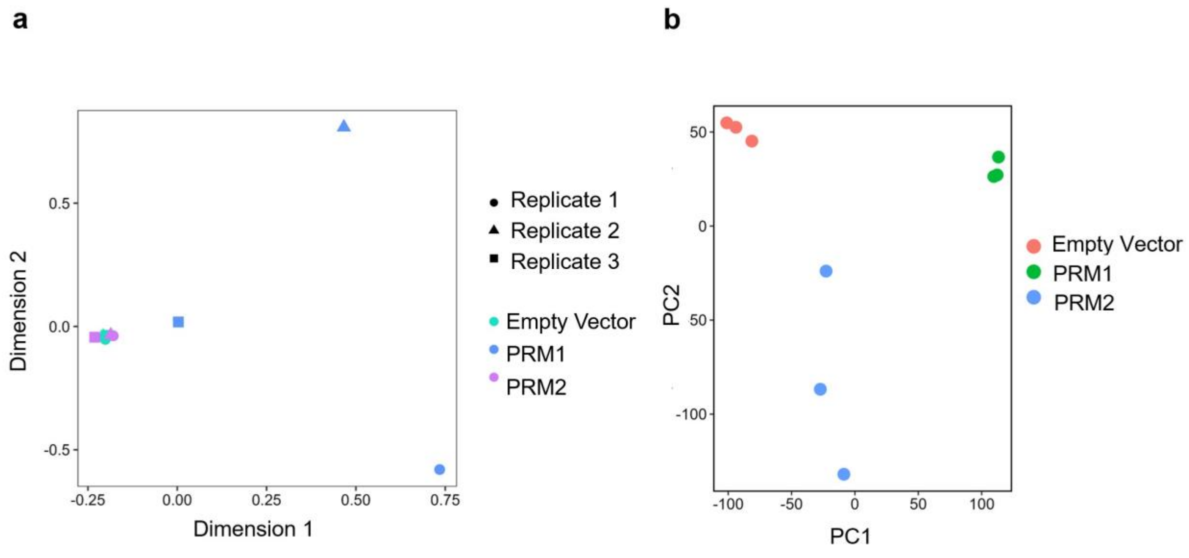


**Figure S3: Sample distribution of DNA methylation and RNA-sequencing**
**a**) MDS analysis of the top 10,000 differentially methylated regions in control and PRM1 and PRM2-transfected cells **b**) PC analysis of RNAseq from control and PRM1 and PRM2-transfected cells

**Supplemental Table 1: Primers used for qRT-PCR**

| Target gene | Primer Name | Direction | Sequence |
|---|---|---|---|
| GAPDH | GAPDH F | Forward | GAAGGTGAAGGTCGGAGTC |
| GAPDH | GAPDH R | Reverse | GAAGATGGTGATGGGATTTC |
| PRM1 (human) | hPRM1f1 | Forward | ATCCACCAAACTCCTGCCTG |
| PRM1 (human) | hPRM1r1 | Reverse | ACAGGCGGCATTGTTCCTTA |
| PRM2 (human) | hPRM2f1 | Forward | CGTCGAGGTCTACGAGAGGA |
| PRM2 (human) | hPRM2r1 | Reverse | CCTGGTTCTGCAGCCTCTG |
| Prm1 (mouse) | mPrm1f1 | Forward | CCGCCGCTCATACACCATAA |
| Prm1 (mouse) | mPrm1r1 | Reverse | GTTTTTCATCGGACGGTGGC |
| Prm2 (mouse) | mPrm2f4 | Forward | TATGGGAGGACACACAGGGG |
| Prm2 (mouse) | mPrm2r4 | Reverse | CCTCCTTCGGGATCTTCTGC |

**Supplemental Table 2: Antibodies used in this study**

| Target | Host Species | Manufacturer | Catalog Number | Concentration for immunostaining | Concentration for Western blot |
|---|---|---|---|---|---|
| **Primary Antibodies** | | | | | |
| H3K4me1 | Rabbit | Abcam | ab8895 | 0.5 μg/ml | 1:3000 |
| H3K9me3 | Rabbit | Abcam | ab8898 | 0.5 μg/ml | 1:2500 |
| H3K27ac | Rabbit | Abcam | ab4729 | 1 μg/ml | 1:3000 |
| H3 | Rabbit | Abcam | ab1791 | | 1:3000 |
| **Secondary Antibody** | | | | | |
| Goat anti Rabbit Alexa Fluor 594 | | Invitrogen | A11012 | 2 μg/ml | |
